# Exploring an experiment-split method to estimate the generalization ability in new data: DeepKme as an example

**DOI:** 10.1101/2021.03.19.436140

**Authors:** Guoyang Zou, Lei Li

**Affiliations:** School of BasicMedical Science, Qingdao University, China; School of Basic Medicine, Qingdao University, Qingdao 266071, China; School of Data Science and Software Engineering, Qingdao University, Qingdao 266071, China; Qingdao Cancer Institute, Qingdao University, Qingdao 266071, China

**Author notes:** Corresponding author; ^1^, ^2^.

## Abstract

A Large number of predictors have been built based on different data sets for predicting different post-translational modification sites. However, limited to our knowledge, most of them gave an overfitting estimation of their generalization ability in new data because of the intrinsic trait—not considering the experimental sources of the new data—of the cross-validation method. Thus, we proposed and explored a new method—the experiment-split method—imitating the blinded assessment to deal with the overfitting problem in the new data. The experiment-split method logically split the training and test data based on the data’s different experimental sources, and the new data can be regarded as the data from different experimental sources. To specifically illustrate the experiment-split method, we combined an actual application, DeepKme—a predictor built by us for the lysine methylation sites, to demonstrate how it be used in the true scenarios. We compared the cross-validation method with the experiment-split method. The result suggested the experiment-split method could effectively relieve the overfitting compared with the cross-validation method and may be widely used in the field of identification participated by multiple experiments. We believe DeepKme would facilitate the related researchers’ deep thought of the experiment-split method and the overfitting phenomenon, and of course, advance the study of the lysine methylation and similar fields.

## Introduction

The central dogma is not able to determine the phenotype of organisms exclusively, and the environment has been always playing an important role in the growth and development of life forms. To respond to the complex and multiple external stimulations, the protein as the main performer of biological functions must flexible enough to quickly behave different morphological structures and thus functions. Compared to synthesizing a new protein, adding a molecular on the original protein to change its structure and function, which is called post-translational modification(PTM), would be much faster to respond to the outside stimulation, making the life more chance surviving [1]. As one of the most important PTMs, the history of the lysine methylation of proteins was winding that the first lysine methylation phenomenon was found on bacterial flagellar proteins, then on mammalian histones, then on nonhistones, and some enzymes such as protein lysine methyltransferases (PKMTs) and demethylases (KDMs) were also be found gradually, meaning the process of lysine methylation was a dynamic and reversible, and so they may play an important role in cell signaling and regulating [2]. Recent studies have been accumulating more and more evidence showing the important interaction between lysine methylation and cancer and thus cancer therapy [3–5]. However, usually, the important first step is to determine the precise position of the lysine methylation, and other steps would be after that.

Nearly all of the lysine methylation sites were identified through high-throughput mass spectrometry(MS) methods combining with enriching the objective peptides or proteins ahead of time through isolating approaches such as affinity purification, immunoprecipitation, etc. Differing from the experiment-extensive and time-consuming MS identification, the computational prediction of the lysine methylation sites would overcome all the defects and keep going along the hypothesis-driven scientific way to push science forward. Limited to our knowledge, the first predictor including the prediction for lysine methylation is built by Daily et al [6] in 2005 using the intrinsic disorder information as the feature and support vector machine (SVM) as the learning model. A predictor called AutoMotif [7] in 2005 used the linear functional motif (LFM) as the feature and SVM as the learning model. Machine learning models are powerful tools of classification, and SVM is one of the most popular tools. A predictor called Memo in 2006 also used SVM as its learning model. The above are early predictors, which were too flexible to form a fixed pattern in building predictors. Afterward, some relatively fixed patterns in building predictors came into beings, such as the data collection based on sequence information with all other information inferred from it, feature engineering, traditional machine learning methods, performance evaluation, and comparison [8–24]. Recently, as the increasingly popular of deep learning which using a different structured deep artificial neural network as the learning method, some predictors [25–27] basing on deep learning also were built for the prediction of lysine methylation sites even if may not be targeted. However, the fatal shortcoming of the computational prediction is the false-positive proportion is high and has always been far miscalculated because of the unrepresentative test set [26,28–32]. Although there may be predictors which would be vetted very well in the future, some experimentalists would more hope to get actionable predictor right away having been strictly assessed imitating most of the practical rather than investigate many predictors by themselves which would be very troublesome because of the big variation through the data sets, modeling methods, programming tools, and availabilities. The core of the overfitting phenomenon is the construction method of the test set, and a traditional way is the cross-validation method. Limited to our knowledge, most of the predictors gave an overfitting estimation of their generalization ability in new data [29,30]. The immediate cause, the objective reason, the subjective reason, and the root causes are as follows.

1. The immediate cause is the test set can not unbiasedly represent the new data, in other words, the test set and the new data have a different distribution.
2. The objective reason is any old data can not unbiasedly represent the new data.
3. The subjective reason is the developers hope and believe the cross-validation method can get the most accurate estimation of the generalization ability in new data, and an important side is the overfitting happens even in their own test set because of the inadvertent mistreatment of the test set as the validation set.
4. The root cause is, from the perspective of cognosciblism, our recognization of any world being through any approach is inevitably biased.

Despite all these, it’s never unallowed to propose some new methods to try to solve the problem. Due to we can’t stop the above objective reason and root cause, we proposed and explored a new method called the experiment-split method trying to solve the problem through changing our subjective factors—the cross-validation which only considers the general key distinguishing a sample from the others. In the experiment-split method, we used the higher-level key—the experimental source—to split our data, and we put the data from one experimental source each time on the test set and the other data on the training set, while the cross-validation is that after splitting the data into several the same-size pieces, the data was put from one piece each time on the test set and the other data on the training set. The two methods were compared on our new proposed predictor called DeepKme for lysine methylation sites. And the roles of DeepKme are as follows.

1. First, DeepKme was used to compare the experiment-split method with the cross-validation method.
2. Second, only by building a reliable—not overfitting—predictor through the experiment-split method can the experiment-split method achieve its aim, thus we should make DeepKme a usable and reliable predictor receiving the practical examination.

Thus, we used a different pattern from before to complete the predictor, the differences are as follows:

### 1 A different construction of testing negative sites from before

Most of the researchers used the lysine sites not identified methylated on methylated proteins as negative sites, but we use the lysine sites on proteins not identified methylated as negative sites. The known human proteins have about 700,000 lysine sites with only 5000 around being identified methylated. We randomly chose 20,000 sites from the left 650,000 sites as negative sites for training, and another 20,000 sites for conservative testing [32]. To evaluate the model performance in positive sites, we split the positive sites into 10 pieces, and use 10 fold cross-validation to reduce the random error in testing. But we don’t use 10 fold cross-validation in negative sites because the number 20,000 is big enough meaning stable enough and the consideration of extra computational cost. The operation makes the testing sites more representative because the negative sites in testing sites are conservative in most true scenarios for the existence of false-negative sites which means the true specificity is higher than the specificity in testing sites while the true sensitivity remains the same.

### 2 Two different construction of testing positive sites for comparison, one of which is the same as before and the other is different

We have two evaluating methods for two different practical scenarios, one is the experiment-split method which split the positive sites into different groups by experiment sources, and the training positive sites must have different experiment sources from testing positive sites, the other is the traditional cross-validation method.

### 3 Multi-task deep learning model

As the lysine methylation has three or more degrees namely mono-, di- and tri-methylation, and the combination modes are a little too many so that it’s a low effective way to build one model for each mode, the multi-task learning model can deal with multiple tasks each time and the common layers of different tasks may play a role as regularization to reduce overfitting because it reduces the parameter space of the model.

## Methods

### Datasets

The data about lysine methylation sites were collected from several different sources (Fig 1) (S1 and S2 Tables) but mainly from PhosphoSitePlus in quantity. Then during the data cleaning, some unreliable data was dropped, and followed by data organization with all lysine methylation sites represented in uniformed format (Table 1).

**Table 1.** The meaning of header in the data table.

| Column | Meaning |
| --- | --- |
| Entry | Primary (citable) accession number in UniProt. |
| Position | The position of the lysine in a protein and it starts from 1. |
| SeqWin | The peptide sequence with the lysine in the center and its length is fixed on 61. |
| Evidence | The Pubmed or CST ID corresponding to the site. |
| Sequence | Protein sequence. |

**Fig 1.**
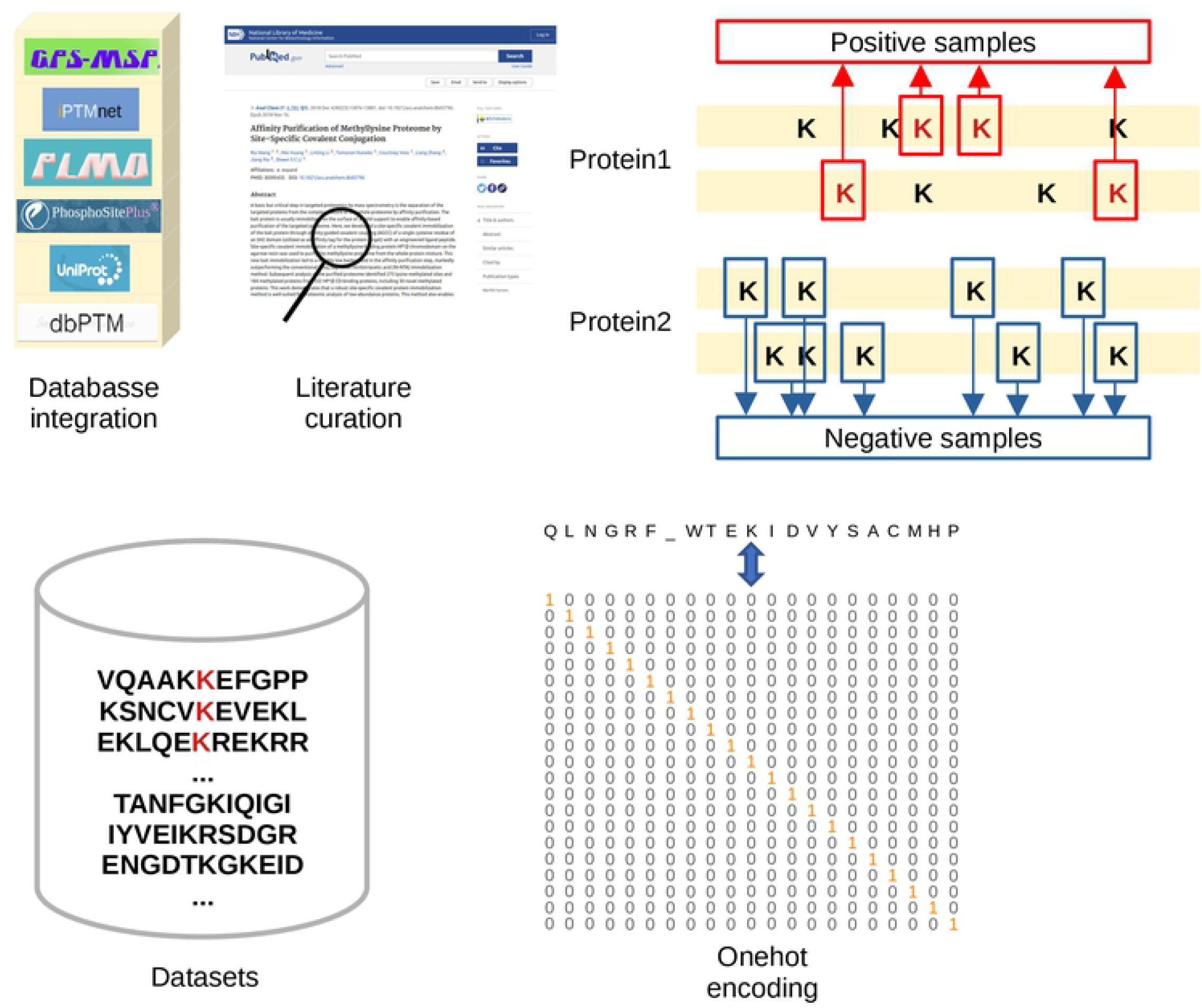
The working flow of data collection.

Similar to the practice of many researchers in the past, we used 61 lengths of sequence windows with “K”(lysine) in the center and other amino acids or the padding “_” representing no amino acid in two sides and corresponding positions in corresponding proteins. The last step was redundancy reducing because the uniformed data obtained in the last step may be duplicated depending on your definition of the unique primary key or id and the duplicated data was meaningless to some extend. In our computational experiment, the key or id was defined to be the sequence window because which is the input of our model, corresponding to only one output in a format of a value or a series of value namely a vector and all the inputs data of the testing set or training set or positive sample set or negative sample set was unique. Note, we did not reduce redundancy below 100% similarity, although many others claimed that the operation could avoid overfitting or improve the performance of the model. If using Entry and Positon as the key of the data table, we can get a summary of the data size from different sources (Table 2). The PHP [33] is a constantly updated database of several common PTMs such as phosphorylation, acetylation, ubiquitination, methylation, etc (Fig 2). Although we collected sites from multiple sources, most sites can be found in PHP and the specific types such as mono-, di-, and tri-methylation are labeled in most of the sites. However, the key consisting of Entry and Position is not suitable for modeling, and using different keys would determine different sample numbers (Table 3) and have a different function (Table 4). So another key consisting of SeqWin is more appropriate to represent the sample set, and other keys can be found in (Table 4).

**Table 2.** A summary of the data size of sites from different sources.

|  | PLMD | MSP | PHP | dbPTM | Paper | iPTMnet | uniprot | All |
| --- | --- | --- | --- | --- | --- | --- | --- | --- |
| Km1 | 0 | 279 | 4252 | 0 | 54 | 0 | 175 | 4423 |
| Km2 | 0 | 107 | 529 | 0 | 53 | 0 | 65 | 635 |
| km3 | 0 | 56 | 326 | 0 | 65 | 0 | 59 | 419 |
| Kme | 1901 | 695 | 5004 | 3431 | 172 | 210 | 228 | 5450 (Key: Entry, Position) |

**Table 3.** The positive sample size for different keys in histone and nonhistone.

|  | In histone | In nonhistone | All |
| --- | --- | --- | --- |
| Protein (Key: Entry) | 45 | 2944 | 2989 |
| Site (Key: Entry, Position) | 221 | 5229 | 5450 |
| Site (Key: SeqWin) | 162 | 5067 | 5229 |

**Table 4.** The function of several keys in the data table.

|  |  |
| --- | --- |
| Key | Function |
| Entry | Distinguish different proteins |
| Entry, Position | Distinguish different sites |
| Evidence | Distinguish the sites from different experiments |
| SeqWin | Distinguish different peptide sequences of sites |
| Label | Distinguish known positive or unknown sites |

**Fig 2.**
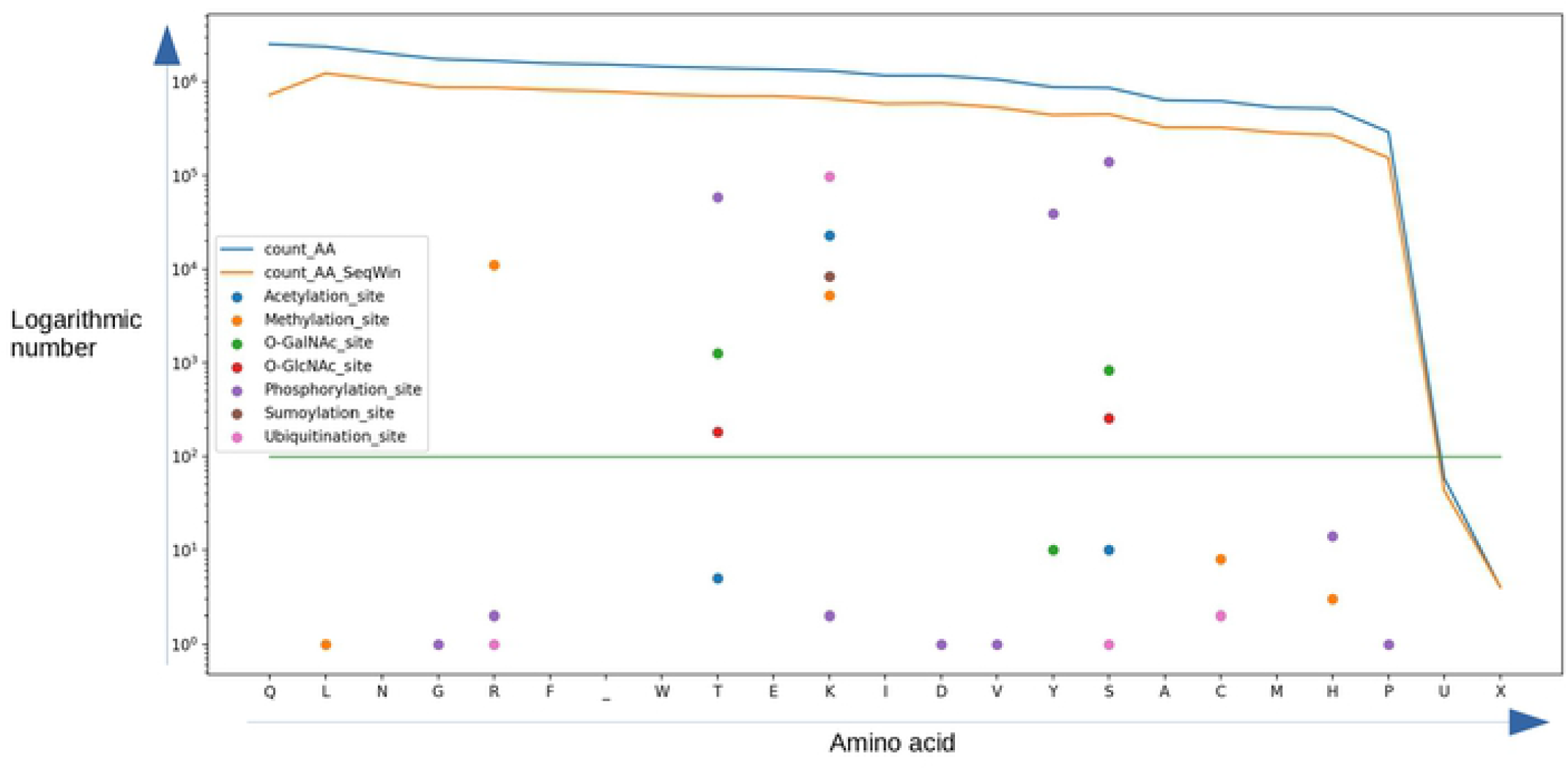
The numeric summary of several PTM sites in human proteome from PHP. The horizontal axis represents the 20 amino acids and a padding “_”. The vertical axis represents the logarithmic number of each PTM. The “count_AA” represents the number of each amino acid in the human proteome. The “count_AA_SeqWin” represents the number of sequence windows of each amino acid site.

### Positive and Negative samples

#### Positive samples

There is not any doubt that all the known lysine methylation sites were used as positive samples because of a very little false positive. Note, the evidence information about them was kept.

#### Negative samples

Because the negative samples were not identified with high specificity, the only way is to choose those unidentified sites as the negative samples. Differing with others selecting the unidentified sites in identified proteins assuming that these so-called negative samples have very few false-negative proportions, we choose the unidentified sites as negative samples in unidentified proteins with another assumption that these so-called negative samples still have very few false-negative proportions but are more representative for most scenarios because that the unidentified proteins are far more than the identified proteins and from proportion, most scenarios are aimed at unidentified proteins. We randomly choose 40,000 sites using SeqWin as key from about 638,805 negative sites for the negative samples for modeling (Table 5) (S5 Table).

**Table 5.**
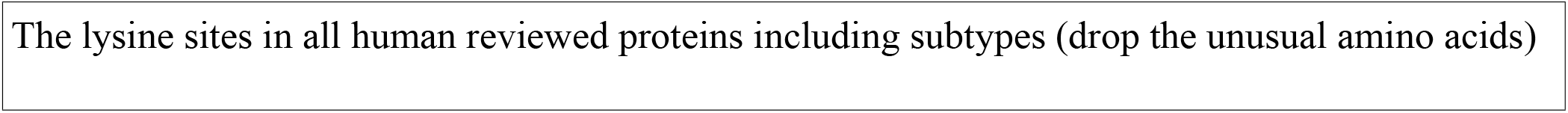
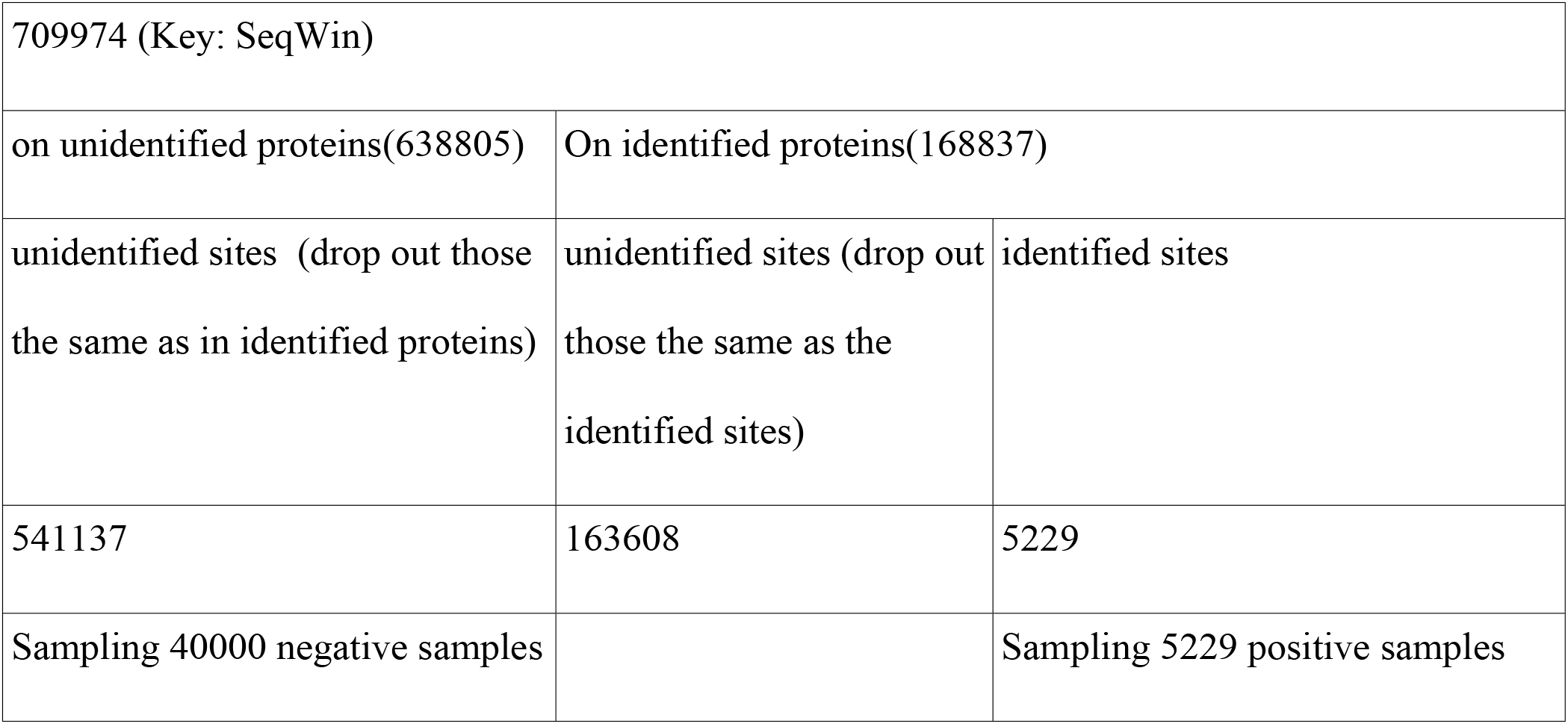
The overview of the lysine methylation sites in the human proteome.

|  |
| --- |
| The lysine sites in all human reviewed proteins including subtypes (drop the unusual amino acids) |

|  |  |  |
| --- | --- | --- |
| 709974 (Key: SeqWin) |  |  |
| on unidentified proteins(638805) | On identified proteins(168837) |  |
| unidentified sites (drop out those the same as in identified proteins) | unidentified sites (drop out those the same as the identified sites) | identified sites |
| 541137 | 163608 | 5229 |
| Sampling 40000 negative samples |  | Sampling 5229 positive samples |

### Testing set and training set

The splitting approach of the testing set and training set in positive samples have two ways, one is the traditional method, the other is the experiment-split method.

### For the traditional method

First, we should know the duality of the meaning of cross-validation (Tables 6, 7), then we used the 10 fold cross-validation method in the second meaning, but there were some differences:

**Table 6.** The duality of cross-validation.

| Cross-validation | Meaning | Ability |
| --- | --- | --- |
| First meaning | To more stably and comprehensively evaluate the model's performance using all data as both testing set and training set, but at each time, they are mutually exclusive. | The model is not the best, but accurately and objectively evaluated. |
| Second meaning | To more stably and comprehensively adjust the model's hyperparameters to select the best model. | The model is the most promising, but overfitting. |

**Table 7.**
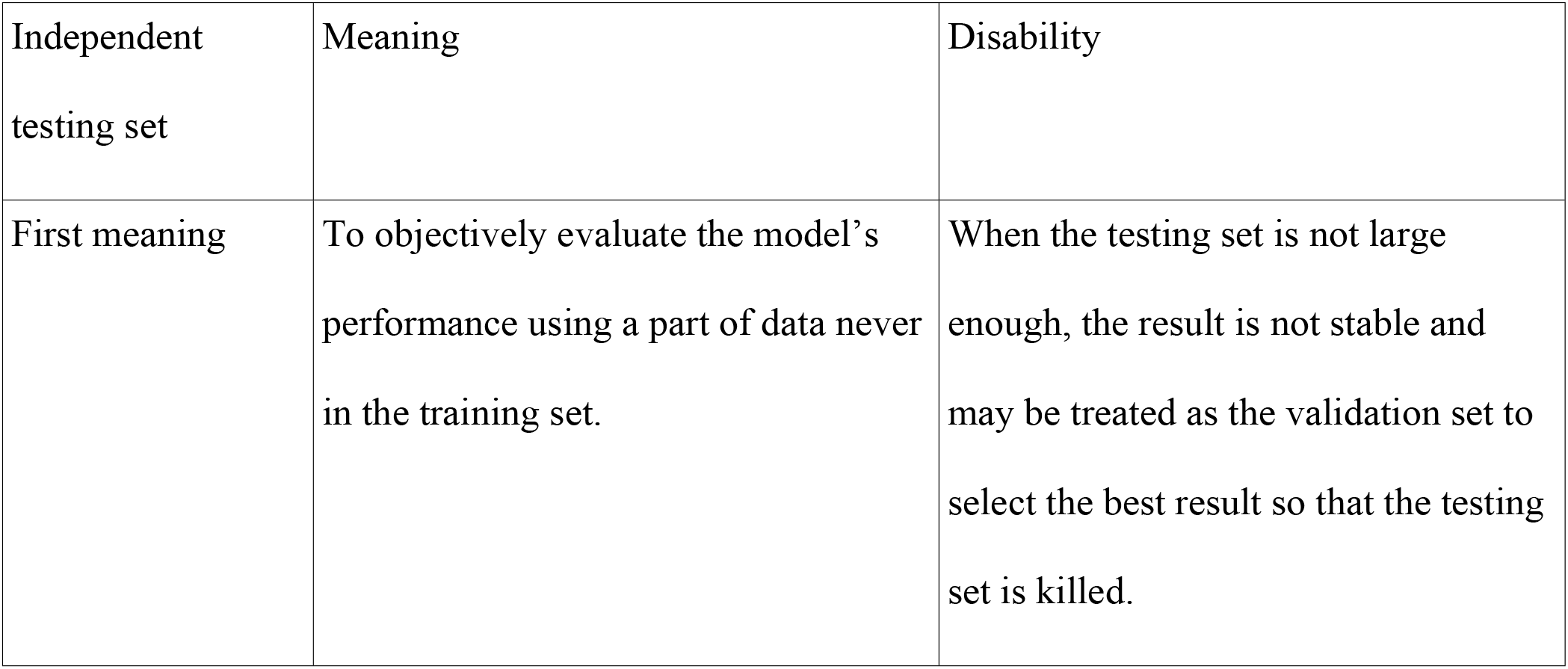
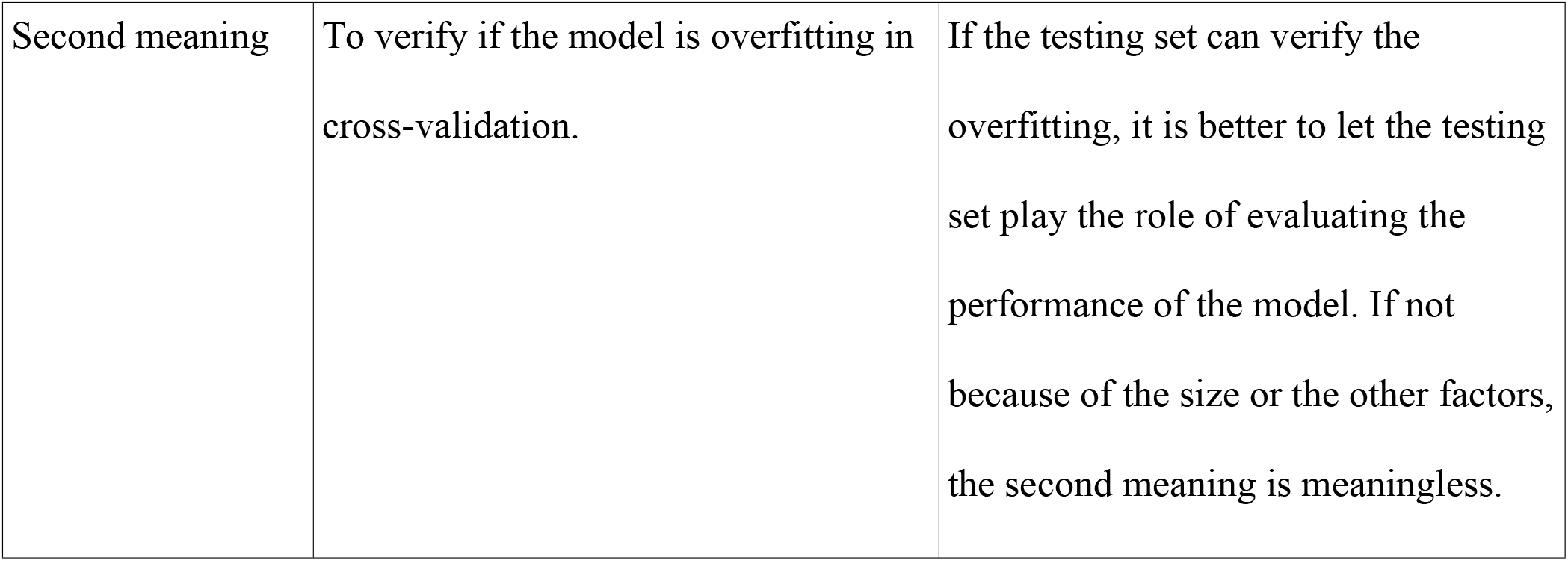
The duality of the independent testing set.

1. Our 10 fold independent testing sets played the role of the validation set, while in the training set, we split out 10 fold validation set for early stopping—when the model’s loss in the validation set didn’t decrease, we stopped training, and choose the minimum loss model as a trained model.
2. The hyperparameters of the multi-task deep learning model were adjusted only on the nine percent—nine-tenths by one-tenth, which were the ratio of the training set in the whole data set and the validation set in the training set, respectively—of the whole data samples. These hyperparameters included the number of convolutional layers, the kernel size, padding or not, the activation function, the max-pooling size, the dropout rate, batch normalization or not, the filter number, and so on. The model’s power contradicts the accurate evaluation of the model’s power, so we use only a little part of the data set to choose a well-performed model to minimize the overfitting.

### For the experiment-split method

It can imitate the blinded assessment which is thought as perhaps the gold standard evaluating the performance of a predictor, where the experiment-split testing method is described as follows:

1. Each lysine methylation site corresponded to an experiment from which the site was identified, and we use the PUBMED or CST(cell signaling technology) id to represent the experiment source. To represent the practical scenarios, the experiment sources of testing data should be different from that of the training data. So we split out the testing data from all data by experiment sources, and the training data from the rest, obtaining many training-testing data sets with the number equaling to the number of experiment sources.
2. Because different experiment sources may get the same sites, to get a more conservative result, we randomly dropped out the overlapping sites from the testing data and training data (S5 Table).

### The splitting approach of the testing set and training set in negative samples have only one

We randomly chose 20,000 sites from the 638,805 negative sites as the training set, and the other 20,000 sites as the testing set. We don’t use 10 fold cross-validation in negative sites is because the number 20,000 is big enough meaning stable enough and the consideration of extra computational cost. Details are summarized in Table 8.

**Table 8.** The positive and negative sample sizes in the training set and testing set.

| Training set |  | Testing set |  |
| --- | --- | --- | --- |
| positive | negative | positive | negative |
| 4698 | 20000 | 522 | 20000 |
| 10 fold cross-validation (the second meaning) in the inner loop |  |  |  |
| 10 fold cross-validation (the first meaning) in the outside loop |  |  |  |

### Data encoding

Usually, the sample information should be appropriately represented as a number or number sequence namely a vector as the starting point of computation. Here, we use one-hot encoding to represent original site sequence window information, for example, “AKS” as “100,010,001”, and “SKA” as “001,010,100”. Although the simplest way to represent a sequence may be using a unique integer to represent a unique word in the sequence, an evident drawback is these integers have size relationships but word not, causing difficulties to the learning process of the machine learning model. By contrast, the one-hot encoding solves this problem and is often used for sequence representation. Each word of a sequence is represented as a vector with the length number of categories, and the value in each position of the vector corresponds to the existence of a word, then each time when using the vector to represent a word there must be only one position in the vector matches the word, so the number in that position is set to 1, and others are set to 0 (Fig 3).

**Fig 3.**
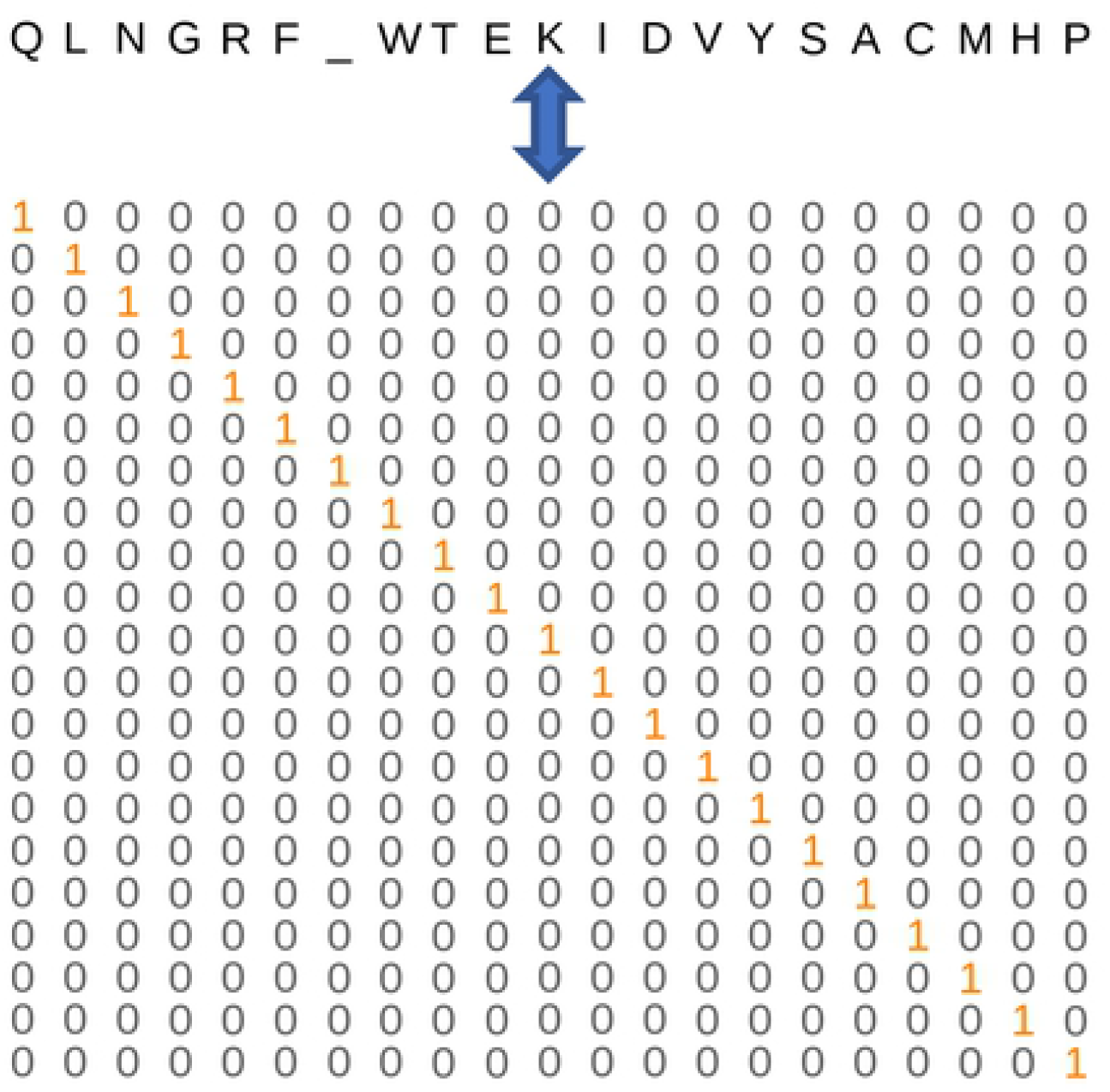
Onehot encoding.

### Model

A neural network has far better fitting power than traditional machine learning models such as SVM, Random forest, and so on. Not very strictly, it can fit nearly any function relation so that it is easy to make the performance in trainset perfect, such as 100% accuracy both in negative and positive samples. However, overfitting in the test set is the main or not strictly the only challenge we face, for instance, perfect performance in trainset but bad in the test set. There are many approaches to alleviate overfitting, such as restricting the weight through regularization to narrow the decision space to a more simple relation basing on Occam’s razor law—Entities should not be multiplied unnecessarily. Here, we use multi-task learning in the computer science field to build our model. On account of the three cases of lysine methylation including mono-, di- and tri-methylation of lysine have different sample volumes with di- and tri-methylation very few samples, a multi-task learning model was applied to share some weights of models for the three cases so that each of them would benefit from it, just like a student learns multiple subjects simultaneously to improve the grade of each subject because some knowledge is easier to be learned from this subject rather than other subjects but can be used to other subjects. An early stopping strategy was applied. The parameters are summarized in Table 9. The graph representation of the model is in Fig 4.

**Table 9.** Parameters of the multi-task deep learning model.

| layer | option | value |
| --- | --- | --- |
| Cov1D | filter number | 256 |
|  | kernel size | 9 |
|  | stride | 1 |
|  | padding | valid |
|  | batchnormalization | yes |
|  | activation | relu |
|  | maxpool | 2 |
|  | dropout | 0.7 |
| Cov1D | filter number | 32 |
|  | kernel size | 7 |
|  | stride | 1 |
|  | padding | valid |
|  | batchnormalization | yes |
|  | activation | relu |
|  | maxpool | 2 |
|  | dropout | 0.5 |
| Dense | size | 128 |
|  | activation | relu |
| Dense (output) | size | 4 |
|  | activation | sigmoid |

**Fig 4.**
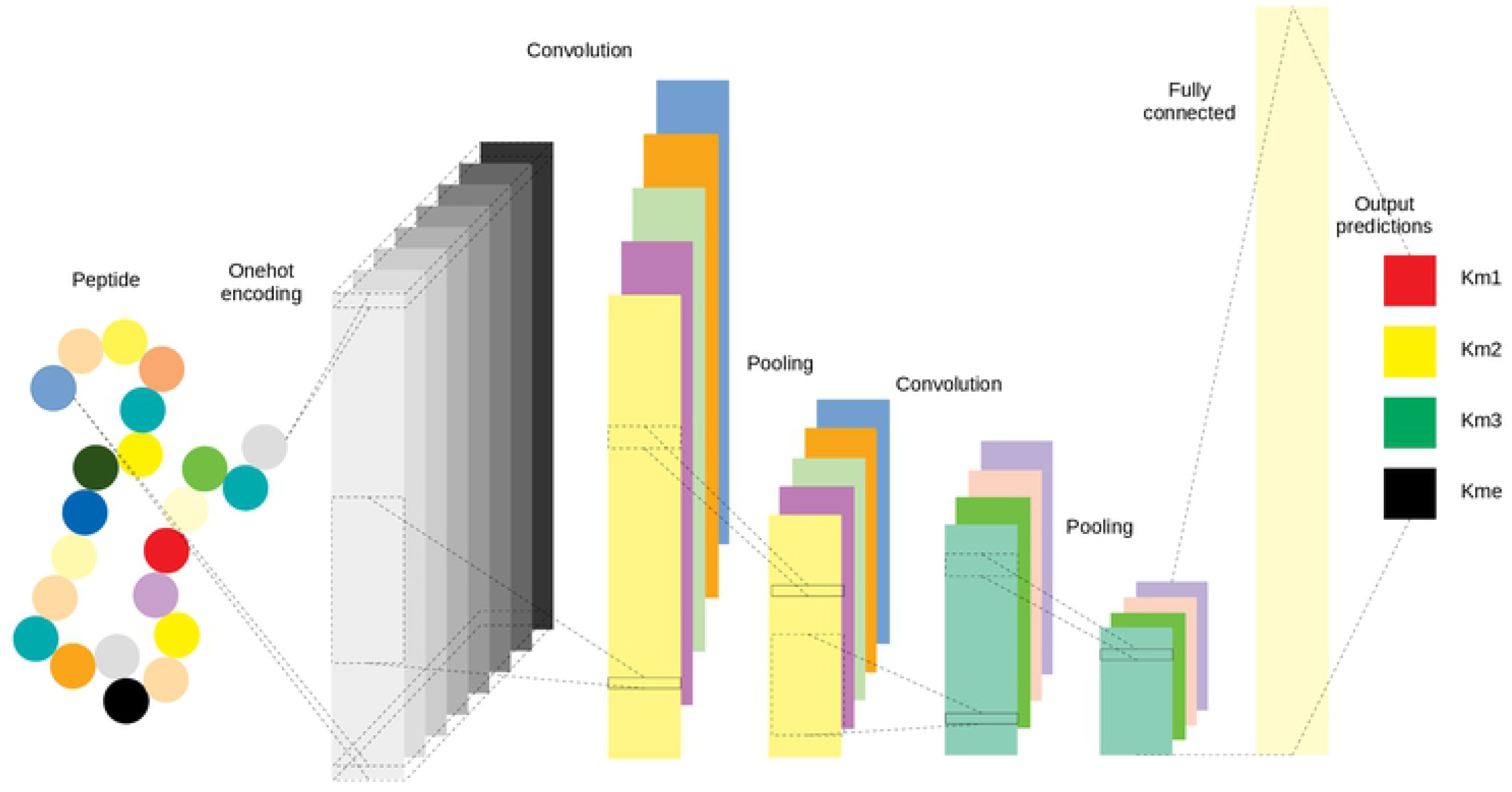
Graph representations of the multi-task deep learning model.

### Comparison

Two comparisons are in Figs 5,6.

**Fig 5.**
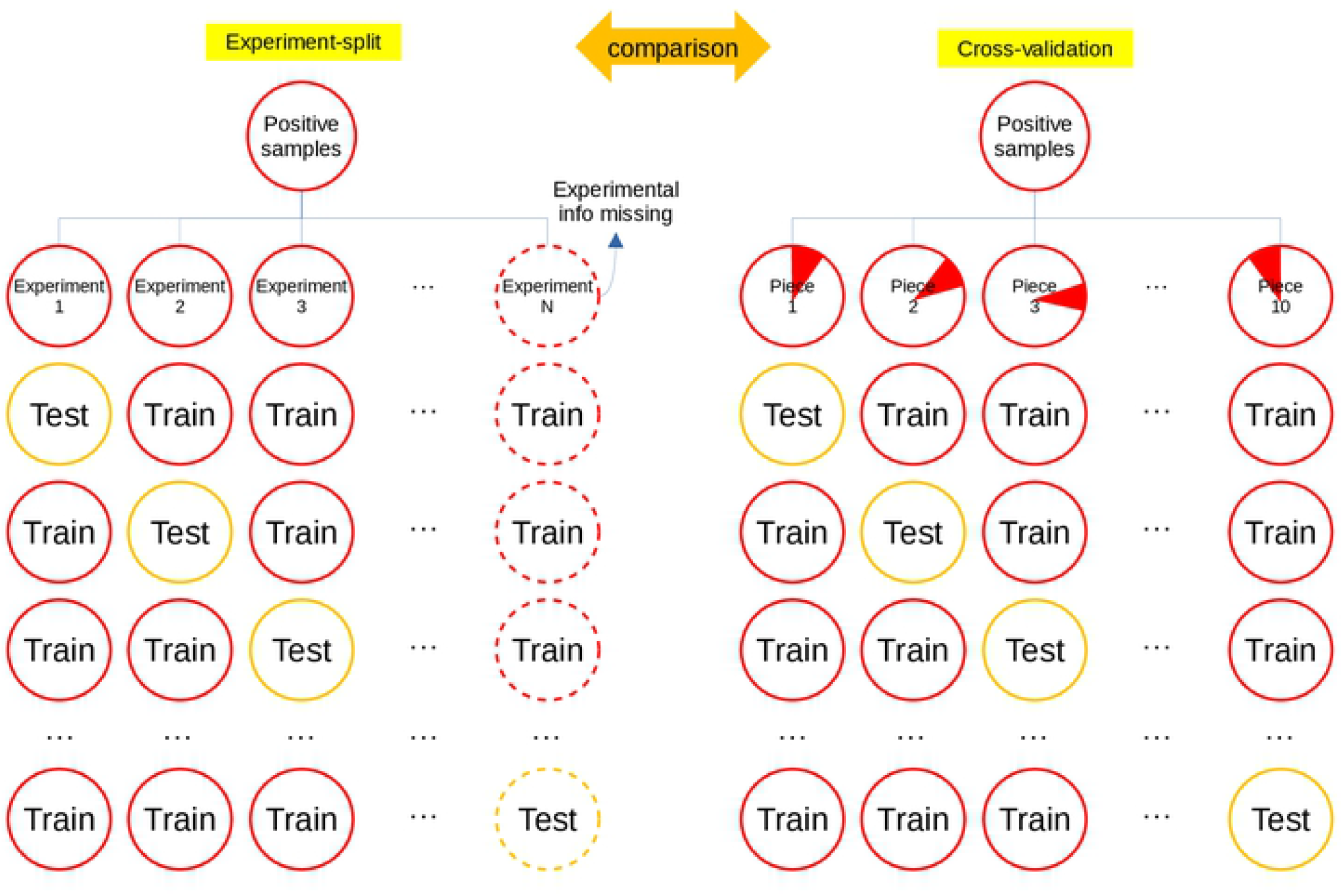
Comparison between the experiment-split method and the cross-validation method.

**Fig 6.**
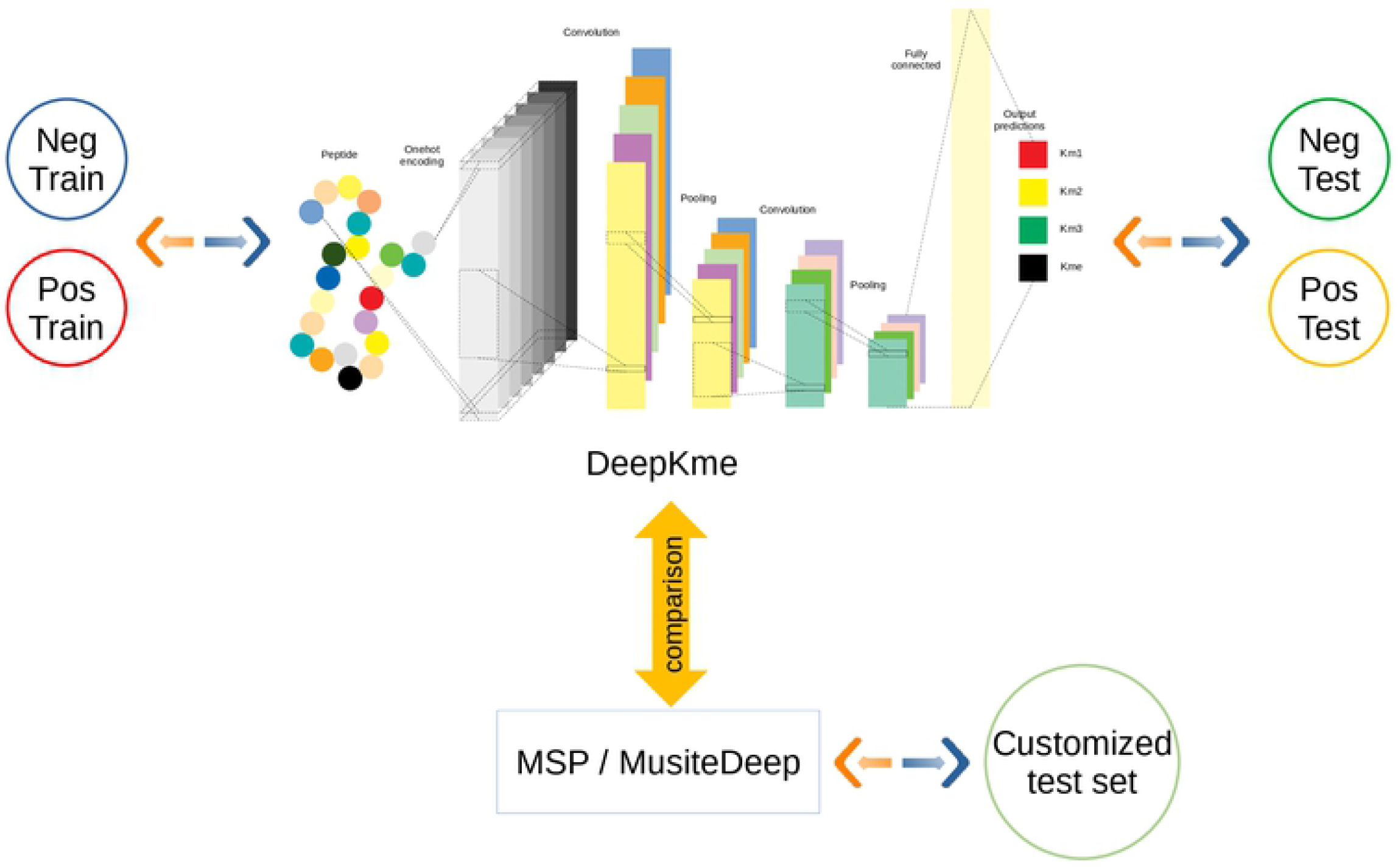
Comparison between DeepKme and the previous predictors(MSP/MusiteDeep).

## Results and Discussions

### For the traditional method

10 multi-task learning models were randomly trained in each fold using the early stopping strategy, and the result showed that the mono-methylation has the best average performance, followed by, in descending order, *-, tri- and di-methylation respectively (Fig 7) (Tables 10, 11 and 12). Note, the data for Km3 was less than Km2, suggesting the features of Km3 were more evident than the features of Km2. In Table 12, when the data size of Km2 was larger than Km3 which was controlled with its size the same with Km1, the mean AUC of Km2 was far lower than Km1 and Km3, suggesting the sequence features of Km2 were harder to be found, just like the Kme which was the mixture of Km1, Km2, and Km3, even if the data size was far larger, due to the dispersion of their features, the mean AUC was not the max, but in the medium. From the intracellular chemical reaction, the Km1 could be followed by Km2 with Km2 PMTKs, which then may be followed by Km3 with Km3 PMTKs. Maybe the medium state of the lysine methylation is less stable than the side state because of the sequence features unless some factors beyond sequence affect it.

**Table 10.**
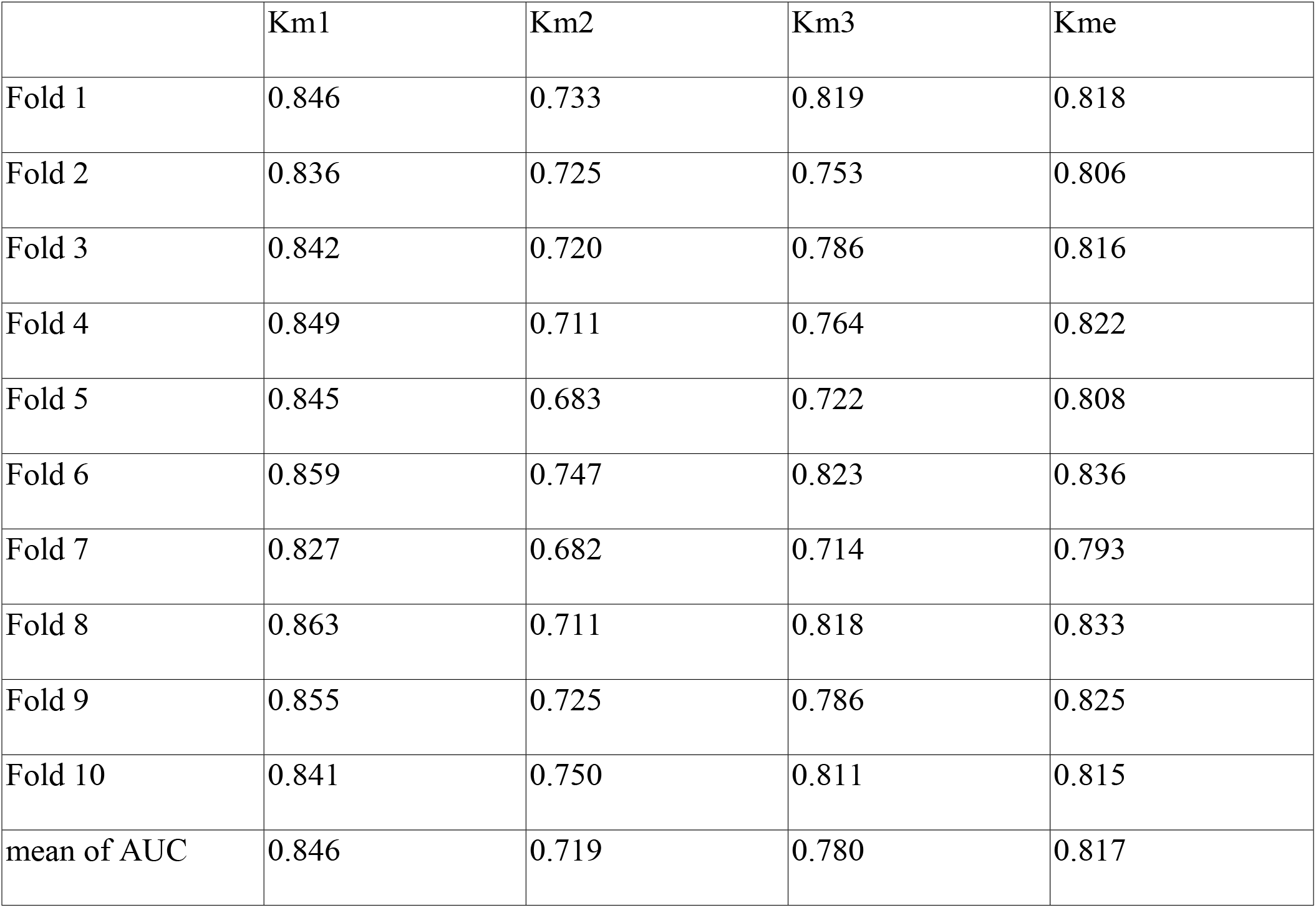
The result of the traditional method using multi-task deep learning model.

**Table 11.** The results of the experiment-split method of different models using the same data set.

|  |  |  |  |  |
| --- | --- | --- | --- | --- |
| Multi-task DL (weighted) | Km1 | Km2 | Km3 | Kme |
| weighted mean of AUC | 0.751 | 0.652 | 0.689 | 0.719 |
| mean of AUC | 0.766 | 0.660 | 0.729 | 0.747 |
| Multi-task RF | Km1 | Km2 | Km3 | Kme |
| weighted mean of AUC | 0.650 | 0.541 | 0.585 | 0.631 |
| mean of AUC | 0.674 | 0.555 | 0.627 | 0.655 |
| Single-task RF | Km1 | Km2 | Km3 | Kme |
| weighted mean of AUC | 0.647 | 0.546 | 0.564 | 0.635 |
| mean of AUC | 0.675 | 0.557 | 0.598 | 0.658 |
| Multi-task RF (weighted) | Km1 | Km2 | Km3 | Kme |
| weighted mean of AUC | 0.672 | 0.548 | 0.579 | 0.639 |
| mean of AUC | 0.693 | 0.552 | 0.587 | 0.677 |

**Table 12.** The result of the traditional method using multi-task deep learning model trained on fewer data.

|  | AUC |  |  |  | Site Number |  |  |  |
| --- | --- | --- | --- | --- | --- | --- | --- | --- |
|  | Km1 | Km2 | Km3 | Kme | Km1 | Km2 | Km3 | Kme |
| Fold 1 | 0.77 | 0.675 | 0.781 | 0.718 | 300 | 460 | 299 | 1165 |
| Fold 2 | 0.768 | 0.666 | 0.708 | 0.713 | 300 | 465 | 304 | 1153 |
| Fold 3 | 0.769 | 0.722 | 0.791 | 0.737 | 300 | 464 | 299 | 1158 |
| Fold 4 | 0.79 | 0.665 | 0.75 | 0.736 | 300 | 462 | 314 | 1174 |
| Fold 5 | 0.771 | 0.699 | 0.722 | 0.713 | 300 | 457 | 307 | 1161 |
| Fold 6 | 0.767 | 0.711 | 0.825 | 0.725 | 300 | 480 | 307 | 1183 |
| Fold 7 | 0.756 | 0.688 | 0.718 | 0.708 | 300 | 466 | 300 | 1160 |
| Fold 8 | 0.782 | 0.684 | 0.766 | 0.736 | 300 | 460 | 297 | 1151 |
| Fold 9 | 0.778 | 0.701 | 0.807 | 0.735 | 300 | 464 | 296 | 1151 |
| Fold 10 | 0.75 | 0.722 | 0.747 | 0.726 | 300 | 470 | 303 | 1175 |
| Mean | 0.77 | 0.693 | 0.762 | 0.725 | 300 | 464.8 | 302.6 | 1163.1 |

**Fig 7.**
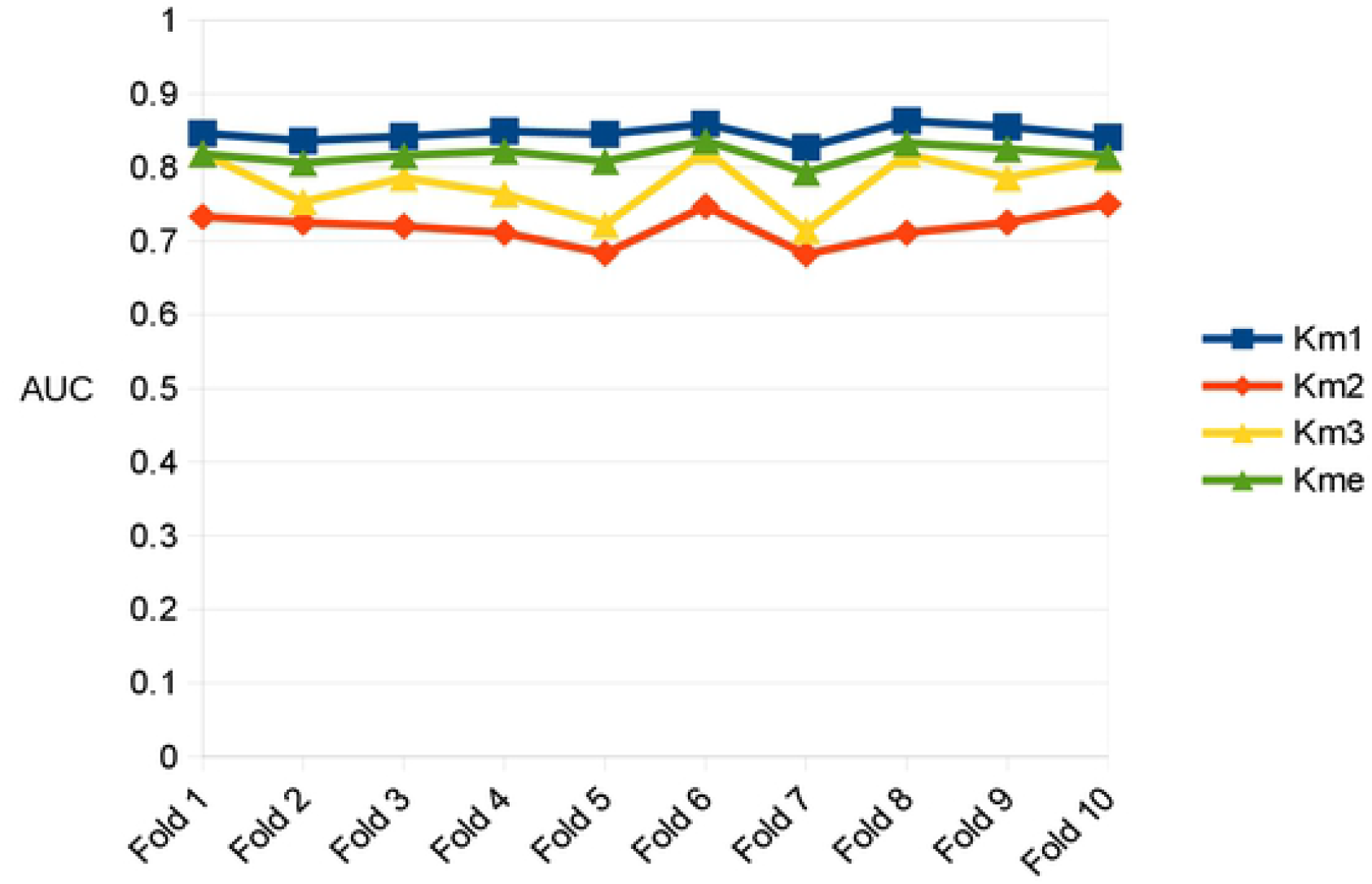
The result of the traditional method using multi-task deep learning model.

### For the experiment-split method

Using the deep learning model, we tested the performance in each experiment-split testing data set, the average AUC is 0.766, 0.660, 0.729, and 0.747 for lysine mono-, di-, tri-, and *-methylation (* represent any of mono-, di- and tri-) respectively (Table 11) (S6 Table). Considering the imbalance of different testing set sizes, we used sizes as weight and weighted average AUC is 0.751, 0.652, 0.689, and 0.719 for lysine mono-, di-, tri-, and *-methylation respectively (Table 11). By contrast, the same model was used in the traditional testing method and the average AUC is 0.846, 0.719, 0.780, and 0.817 for lysine mono-, di-, tri-, and *-methylation respectively (Table 10). Besides, we got nearly the same AUC rank of different experiments in the experiment-split testing method using different models such as RF, indicating that the rank was related mainly to the data itself rather than the model (Fig 8) (Table 11) (S6 and S8 Tables). The previous predictor MSP and MusiteDeep were used for comparison (Fig 9) (Tables 11) (S6 and S7 Tables).

**Fig 8.**
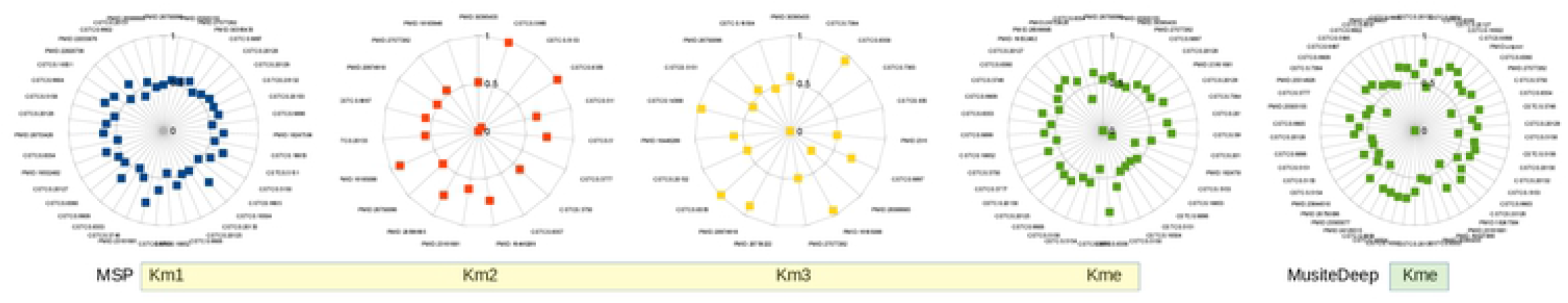
The comparison of multi-task deep learning model and random forest model in the experiment-split testing set. The smaller square is the random forest model. The greater square is the multi-task deep learning model.

**Fig 9.**
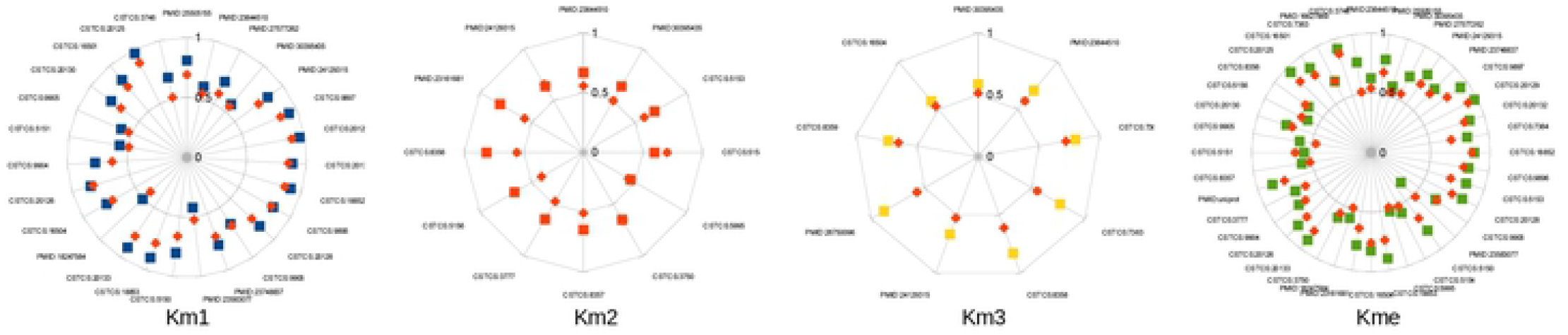
The results of MSP and MusiteDeep in the experiment-split testing set.

We could conclude that:

1. Different model may have different performances in the testing data set, but have nearly the same relative performances in different experiments.
2. If we have better performance in only one or several experiments using the experiment-split method, we would have better performance in other experiments thus better generalization power which is our goal all along. This worth-expecting characteristic is what the traditional independent method does not have, that is, the better performance gained through the traditional independent method does not mean better generalization power.
3. We can use the experiment-split method to construct the benchmark data set, provide a potential approach to solve the difficult problem of benchmarking which plays a very important role in the field of prediction.
4. Each experiment whose number of identified modification sites was greater than a certain number was tested through the experiment-split method and all the metric AUCs of each experiment were plotted in a radar graph which showed the testing performance changed a little much in different experiments. Our model DeepKme has predictability in most scenarios, which was better than MSP and MusiteDeep.

## Supporting information

**S1 Table. The working flow of data collection**.

**S2 Table. The data extraction from UniProt**.

**S3 Table. The URL of each source**.

**S4 Table. The positive and negative sample sizes for sites and protein or other different keys**.

**S5 Table. The summary of sites from different evidence**.

**S6 Table. The result of the experiment-split method using multi-task deep learning model**.

**S7 Table. The result of the experiment-split method using MSP and MusiteDeep**.

**S8 Table. The result of the experiment-split method using random forest model in several settings**.

**S4 File. All the tables of the article**.

